# Comparative RNAi Screens in Isogenic Human Stem Cells Reveal *SMARCA4* as a Differential Regulator

**DOI:** 10.1101/500181

**Authors:** Ceren Güneş, Maciej Paszkowski-Rogacz, Susann Rahmig, Shahryar Khattak, Martin Wermke, Andreas Dahl, Martin Bornhäuser, Claudia Waskow, Frank Buchholz

## Abstract

Large-scale RNAi screens are a powerful approach to identify functions of genes in a cell-type specific manner. For model organisms, genetically identical (isogenic) cells from different cell-types are readily available, making comparative studies meaningful. For humans, however, screening isogenic cells is not straightforward. Here, we show that RNAi screens are possible in genetically identical human stem cells, employing induced pluripotent stem cell as intermediates. The screens revealed *SMARCA4* (SWI/SNF-related matrix-associated actin-dependent regulator of chromatin subfamily A member 4) as a stemness regulator, while balancing differentiation distinctively for each cell type. *SMARCA4* knockdown in hematopoietic stem progenitor cells (HSPC) caused impaired self-renewal *in-vitro* and *in-vivo* with skewed myeloid differentiation; whereas in neural stem cells (NSC), it impaired selfrenewal while biasing differentiation towards neural lineage, through combinatorial SWI/SNF subunit assembly. Our findings pose a powerful approach for deciphering human stem cell biology and attribute distinct roles to *SMARCA4* in stem cell maintenance.

## INTRODUCTION

Stem cells have been in the focus of regenerative medicine due to their two key features: self-renewal and differentiation (Daley et al., 2003; Keller, 2005). Great progress has been made in the stem cell field since the discovery of induced pluripotent stem cells (iPSC) in mouse (Takahashi and Yamanaka, 2006), and in human (Takahashi et al., 2007; Yu et al., 2007). This discovery paved numerous paths including disease modeling and derivation and expansion of somatic stem cells such as HSPC (Sugimura et al., 2017) or NSC (Reinhardt et al., 2013). Although both HSPC and NSC are among the most extensively studied human adult stem cells, it has been challenging to understand the molecular basis behind self-renewal and differentiation, in particular in a comparative way. Interestingly, genes and pathways involved in the fine-tuning of self-renewal and differentiation are often shown to be key players in tumorigenesis (Orkin and Zon, 2008), which renders stem cell research crucial and indispensable in multiple aspects.

Development of RNA interference (RNAi) technology has transformed the pace of functional genetics and provided tools for genome-wide screens. Importantly, RNAi screens have also been performed on mammalian stem cell lines (Ding et al., 2011; Elling and Penninger, 2014; Moffat and Sabatini, 2006) and on patient-derived cells (Camgoz et al., 2018; Wermke et al., 2015) and can be customized for the purpose of each study. Several studies have dissected genetic regulations in human HSPC by lentiviral short hairpin RNA (shRNA) libraries. For instance, *STK38* (Ali et al., 2009), *MAPK14* (Baudet et al., 2012), and cohesin genes (Galeev et al., 2016) have been identified as modifiers of HSPC self-renewal and differentiation. In contrast, NSCs have not been studied in this context, despite being among the most widely studied adult stem cells. Moreover, no comparative study to our knowledge has been performed to identify which genes or regulators function in common, or in a cell type-specific manner in these stem cells.

Ideally, comparative RNAi screens on human stem cells should be performed with isogenic cells, as only isogenic cells can provide an unbiased view for comparative analyses. To address the differences between multiple stem cells that are genetically identical, we hypothesized that cell fate determination is regulated by epigenetic factors. To this end, we chose to study HSPC and NSC, by using iPSC as a bridging-cell type and screened these stem cells with the identical shRNA library targeting 538 epigenetic factors. We identified *SMARCA4*, a chromatin remodeler (Peterson and Tamkun, 1995), as a differential regulator of selfrenewal/differentiation with cell type-specific functions.

## RESULTS

### Isogenic human stem cell derivation via iPSC

To be able to perform comparative RNAi screens on human stem cells, we derived isogenic human stem cells starting from peripheral blood (PB) of a healthy donor. HSPCs were isolated from PB using MACS-based sorting for CD34-positive cells and these cells were directly used for the RNAi screen. By using iPSC as an intermediate cell-type we derived isogenic NSCs, which all together provided the basis for unbiased screens (Figure 1).

**Figure 1.**
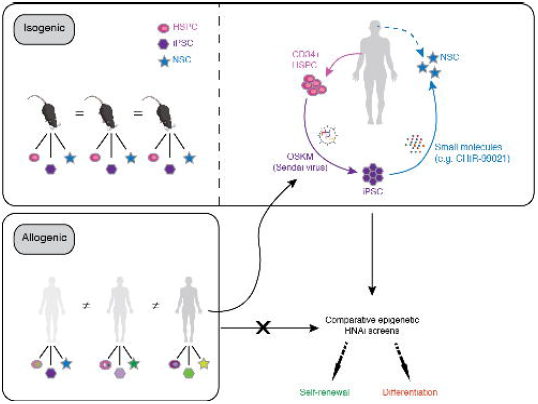
Schematic Representation of Comparative RNAi Screens on Isogenic Human Stem Cells. An alternative approach to derive isogenic stem cells from humans vs. model organisms to perform large-scale screens in a comparative manner. Reprogramming of PB-derived CD34^+^ HSPC to iPSC (bridging cell type) enables differentiation into NSC. Cells derived by reprogramming followed by differentiation serve as isogenic to the starting cell population.

Before performing the HSPC RNAi screen, we tested 7 different suspension culture conditions to find condition boosting CD34^+^ cell proliferation (increasing cell number) while keeping differentiation minimal (CD34^+^ %). We tested conditions, including commercially available small molecules such as pyrimidoindole agonists, i.e. *UM171* (Fares et al., 2014; 2017) and *UM729* (Pabst et al., 2014), as well as cytokines at high concentrations. Addition of UM729 yielded the highest CD34^+^ cell number at minimal differentiation during a 15 day cultivation period (Figure S1). Therefore, we included UM729 for all the following HSPC suspension culture experiments.

As a means of deriving isogenic cell types, we used iPSC, which have been utilized as a source for numerous stem and terminally differentiated cells. While reprogramming HSPC, we opted for a ‘zero-footprint’ method by using the Sendai virus, so that downstream experiments, including RNAi screens and NSC derivation, would not be affected by random genomic integration of the reprogramming factors. We established 2 iPSC lines, which were fully characterized before NSC derivation by iPSC-specific marker expression as well as by the 3 germ-layer differentiation potential (Figure S2). Next, we induced iPSC lines into NSCs by using a cocktail of small molecules (Reinhardt et al., 2013). Loss of pluripotency was confirmed together with the concomitant upregulation of NSC-specific markers. Additionally, similar to the iPSC, we confirmed the functionality of NSCs by differentiation into neurons, astrocytes, and oligodendrocytes (Figure S3). To validate the isogenic nature of the iPSCs and the NSCs, we investigated the isogeneity of these cells by a Short-Tandem Repeat (STR) analysis, which revealed their DNA profiles to be identical to the HSPC population (Table S1). Finally, we performed RNA-seq experiments of the HSPCs, iPSCs and NSCs, to compare their expression profile to published data (Chu et al., 2016; MacRae et al., 2013). As expected, our CD34+ expression profile clustered with two different primary CD34+ expression profiles; iPSC with two embryonic stem cell (ESC) lines; and NSC with two neural progenitor cell (NPC) lines from the literature (Figure S3D). Taken together, we successfully established a minimally invasive approach to derive isogenic human stem cells for unbiased RNAi screens.

### RNAi Screens Identify *SMARCA4* as a Differential Hit

In order to decipher cell fate determinants in isogenic cells, we used a pooled lentiviral shRNA library targeting epigenetic regulators. This library consists of 6482 shRNAs and targets 538 genes - whereby each gene is typically targeted by 12 different shRNAs. As negative controls, 20 non-targeting shRNAs were included (*Renilla* Luciferase, LUC), whereas 6 ribosomal and proteosomal genes served as positive controls (7 shRNAs/gene). We collected the first sample 2 days post transduction (dpt), which served as the baseline for comparison of shRNA representation to later time points. We allowed 5 population doublings between the time points and collected the 2^nd^ time point on 12 dpt, and the 3^rd^ time point sample on 22 dpt. To be able to trace phenotypes back to individual shRNAs, we ensured single shRNA integration by transducing each cell type at low MOI with at least a 150-fold coverage of the library. From each time-point, genomic DNA was isolated from the cells and PCR amplified fragments covering the shRNA sequences were subjected to next generation sequencing (NGS) to identify how shRNA representation changed over time (Figure 2A and Table S2).

**Figure 2.**
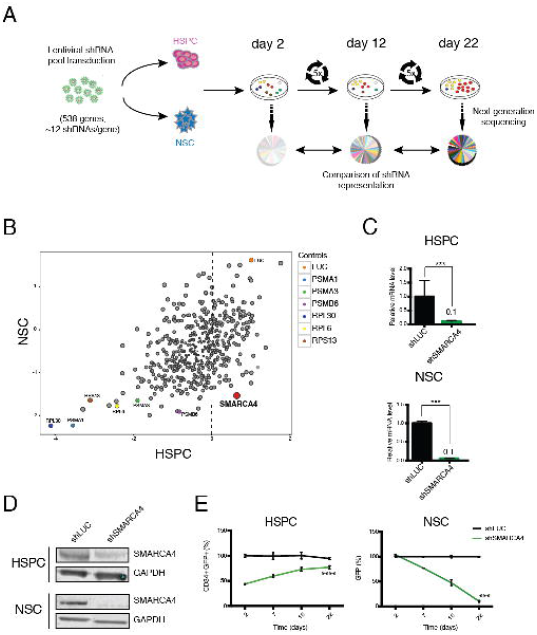
RNAi Screens Identify *SMARCA4* as a Hit with Divergent Phenotypes. (A) Screen strategy and timeline for the pooled shRNA screens. (B) Comparative analysis of the HSPC- and NSC-RNAi screens. Each dot represents a gene with the mean z-score of all targeting shRNAs. *SMARCA4* (enriched in HSPC vs depleted in NSC screen) is depicted in red. Positive controls and the negative control (LUC) are highlighted. See also Table S2. (C) mRNA levels of *SMARCA4* upon knockdown in HSPC (top) and NSC (bottom) measured by qRT-PCR. (D) Confirmation of the *SMARCA4* knockdown at protein level by western blot. GAPDH served as loading control. (E) Validation of screen phenotypes. Enrichment in HSPC based on CD34^+^GFP^+^ expression (left) and depletion in NSC based on GFP^+^ cells (right). Values are normalized to day 2 control sample (HSPC) or shLUC at each time point (NSC). Data are represented as mean ± SEM. ***p < 0.001, ****p < 0.0001. See also Figures S1-3.

The screen results showed that the positive control shRNAs were depleted whereas the negative control shRNAs displayed a relative enrichment phenotype, (compared to the depleted shRNAs), rendering the screen successful. In order to evaluate the screen readout, it is important to be reminded that enrichment of shRNAs of a particular gene can be either due to enhanced self-renewal and cell proliferation or blocked differentiation, and vice versa for a depletion phenotype. Among the hits identified, we decided to investigate *SMARCA4* further, because it displayed a variant phenotype. Indeed, knockdown of *SMARCA4* resulted in enrichment of shRNA reads in HSPC whereas in NSC shRNAs targeting *SMARCA4* were depleted (Figure 2B), suggesting that this gene might have differential roles in these stem cell types.

SMARCA4 is one of the two core enzymatic subunits of the SWI/SNF complex, which is generally associated with 10-12 subunits of BRG1-Associated Factors (BAF). As a core subunit, SMARCA4 has ATPase and helicase activities. The ATPase activity provides the energy via ATP hydrolysis, and the helicase activity unwinds the DNA strands with this energy to alter chromatin accessibility and regulate transcription (Trotter and Archer, 2008). SMARCA4 has been shown to act as a transcriptional activator and repressor, hence, has versatile functions (Attanasio et al., 2014).

Initially, we validated the screen results for the respective cell types by transducing cells with a *SMARCA4* shRNA, which had a z-score of <−2 if depleted, and >+2 if enriched (Table S2). For the validation, we first confirmed the knockdown both at mRNA (~90% knockdown efficiency) (Figure 2C) and protein level (Figure 2D). Then, we took a similar approach as in the screen, and monitored the transduced cells for 22 days, based on the shRNA reporter marker expression (additionally with CD34 for HSPC). We were able reproduce the screen phenotypes, where *SMARCA4* shRNA transduced cells were enriched over time in HSPCs, whereas they were depleted in NSCs (Figure 2E). These data provided evidence that *SMARCA4* is a valid hit from the pooled RNAi screens.

### *SMARCA4* Knockdown Impairs HSPC Self-renewal and Skews Differentiation

Results from the large-scale screen and the initial validation suggested that knockdown of *SMARCA4* increases the percentage of CD34-positive HSPC cells over time. To validate this finding, we monitored the change in CD34^+^ cell number as well as in the overall GFP percentage upon *SMARCA4* knockdown. We observed that CD34^+^ expression was retained in the knockdown cells, which led to a relative enrichment compared to the control. In addition, not only the CD34^+^ percentage but also the overall cell number increased upon *SMARCA4* knockdown (Figure 3A, left). In addition, we also observed an increase in total GFP %, regardless of CD34 expression (Figure 3A, right). Therefore, we investigated further whether this enrichment is specific to HSPC and whether *SMARCA4* has an effect on selfrenewal. First, we examined HSPC functionality upon knockdown by the Long Term-Cell Initiating Culture (LTC-IC) assay (Petzer et al., 1996). After 5 weeks of co-culturing shRNA transduced HSPC (GFP^+^) with stromal feeder cells, substantially more GFP^+^ cells were observed in the shSMARCA4 transduced cells compared to the control (Figure 3B) and *SMARCA4* knockdown led to higher overall cell numbers (3,5-fold, Figure 3C). This data supports our previous findings and altogether can explain the enrichment phenotype observed in the HSPC screen.

**Figure 3.**
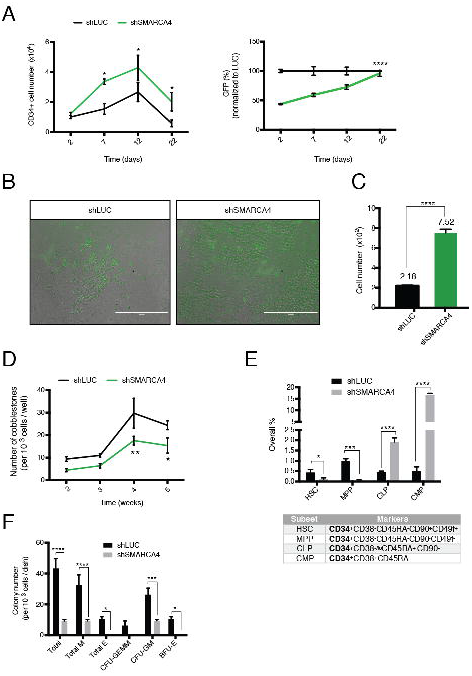
*SMARCA4* Knockdown Phenotypes in HSPC. (A) Increase in CD34^+^ cell number (left) and GFP^+^ percentage (right) upon *SMARCA4* depletion. Values are normalized to day 2 sample control sample (left) or shLUC at each time point (right). Data are represented as mean ± SEM. (B) LTC-IC assay week 5. shRNA transduced HSPC (GFP^+^) cells are shown on the M2-10B4 feeder cells for indicated treatments. Note the increased number of GFP positive cells in the shSMARCA4 treated sample. Scale bar: 200μm. (C) Quantification of the GFP^+^ cell numbers shown in (B). Data from 3 replicates are presented as mean ± SEM. Note the increase in cell number in the shSMARCA4 treated sample. (D) Number of cobblestones in LTC-IC assay between week 2-5. Data from 5 replicates are represented as mean ± SEM. (E) Immunophenotyping of hematopoietic subsets with the indicated marker combination 5 days post shRNA transduction. All subsets analyzed express CD34. Data from 3 replicates are represented as mean ± SEM. HSC: Hematopoietic Stem Cell, MPP: Multipotent Progenitor, CLP: Common Lymphoid Progenitor, CMP: Common Myeloid Progenitor. (F) Colony-unit-forming (CFU) assay. Colonies were imaged 14 days after seeding. Data from 3 replicates are represented as mean ± SEM. M: Macrophage; E: Erythroid; CFU-GEMM: CFU-Granulocyte, Erythrocyte, Macrophage, Megakaryocyte; CFU-GM: CFU-Granulocyte, Macrophage; BFU-E: Burst-Forming-Unit-Erythrocyte. *p < 0.05, **p < 0.01, ***p < 0.001, ****p < 0.0001.

We then evaluated the read-out of the LTC-IC assay by cobblestone formation, which indicates the frequency and activity of HSPC. As of week 2, cobblestone emergence was observed in both shSMARCA4 and control transduced groups. However, despite higher cell numbers in suspension, *SMARCA4* knockdown cells showed a substantial reduction in cobblestone formation (Figure 3D). Thus, *SMARCA4* depletion results in higher CD34^+^ cell number but this increase is not due to enhanced self-renewal of HSPC.

To investigate this observation in more detail, we addressed how *SMARCA4* affects cell differentiation. To this end, we analyzed different hematopoietic subsets in suspension culture, based on immune-phenotypes: HSC, multipotent progenitor (MPP), common-lymphoid (CLP) and -myeloid progenitor (CMP), both derived from MPP (Akashi et al., 2000; Kondo et al., 1997; Notta et al., 2011). Of note, all these cell populations express CD34. Interestingly, *SMARCA4* knockdown resulted in significant reduction of the HSC and MPP populations; whereas both CLP and CMP showed an increase by 4- and 32-fold, respectively (Figure 3E).

To test whether differentiation capacity is altered upon *SMARCA4* knockdown, we performed the Colony-Forming Unit (CFU) assay, where cells are cytokine-induced to form colonies of myeloid origin. Upon 2 weeks of culture, we noted a significant reduction in overall colony numbers. Moreover, *SMARCA4* knockdown samples gave rise only to CFU-GM (granulocyte-macrophage), whereas no erythroid colony formation was observed (Figure 3F). Together, these data suggest that *SMARCA4* depletion in HSPC leads to reduced self-renewal and an imbalanced differentiation towards the myeloid lineage *in vitro.*

### *SMARCA4* Knockdown Impedes *In vivo* Engraftment of HSPC

We next investigated how *SMARCA4* knockdown affects hematopoiesis *in vivo.* To be able to evaluate the change in percentage of the GFP^+^ cells, we transplanted (5×10^5^) HSPCs that were infected at a 30-35% transduction rate with either shSMARCA4 or shLUC into NSGW41 mice (Cosgun et al., 2014) (Figure 4A). To investigate the Long-Term HSCs (LT-HSC), we analyzed mice 6 months after transplantation (Zon, 2008). Upon bone marrow (BM) analysis of GFP^+^ cells we did not observe a significant difference in human chimerism between *SMARCA4* knockdown and control mice (Figure 4B). However, among the engrafted human cells (hCD45^+^), engraftment rate of GFP^+^ cells were strikingly low upon *SMARCA4* knockdown (Figure 4C). These results indicate that *SMARCA4* is essential for LT-HSC maintenance, and consequently for hematopoiesis.

On the one hand, our findings show that *SMARCA4* knockdown leads to exit from self-renewal and impedes engraftment *in vivo*; on the other hand, it directs differentiation to progenitors of both myeloid and lymphoid lineages, however, with less capacity of further myeloid differentiation. Altogether, *SMARCA4* is an important regulator of hematopoiesis as its loss leads to impaired self-renewal together with expansion of CD34^+^ cells and distorted differentiation *in vitro* and impeded engraftment *in vivo*.

**Figure 4.**
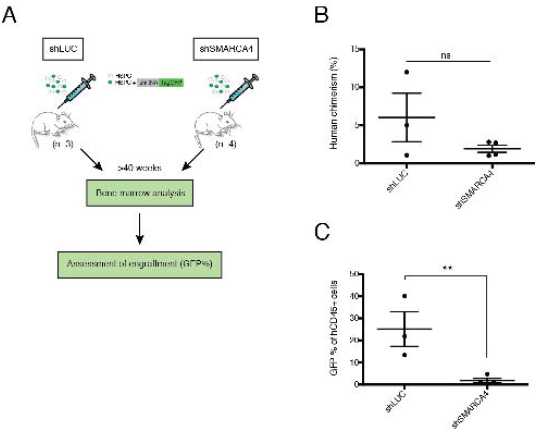
*SMARCA4* Depletion Impairs HSC Engraftment *In vivo*. (A) Schematic representation of the transplantation experiment. NSGW41 mice injected with 5×10^5^ HSPC carrying the control (n=3 mice) and *SMARCA4* shRNA (n=4 mice). Bone marrow was analyzed >40 weeks after transplantation. (B) Human chimerism based on hCD45 expression. (C) Engraftment based on GFP percentage of the hCD45 cells in the control vs *SMARCA4* shRNA transplanted mice. **p < 0.01; ns, not significant.

### Loss of Self-renewal and Cell Detachment in NSC

In contrast to the enrichment of shRNAs targeting *SMARCA4* in the HSC screen, the same shRNAs were depleted in the NSC screen (Fig. 2B). To validate the screen phenotype in NSC, we transduced NSCs with lentiviral particles expressing *SMARCA4* shRNAs in conjunction with GFP and monitored the cells over time by fluorescence microscopy. Strikingly, we observed a very drastic change in morphology of the transduced cells, where *SMARCA4* knocked-down cells budded off (Figure 5A, i) and detached, followed by formation of spheres floating in suspension (Figure 5A, ii). We confirmed this phenotype alternatively by chemically inhibiting SMARCA4 at its bromodomain, using an inhibitor, namely PFI-3 (Gerstenberger et al., 2016; Vangamudi et al., 2015), which reproduced the shSMARCA4 knockdown morphology (Figure 5B).

**Figure 5.**
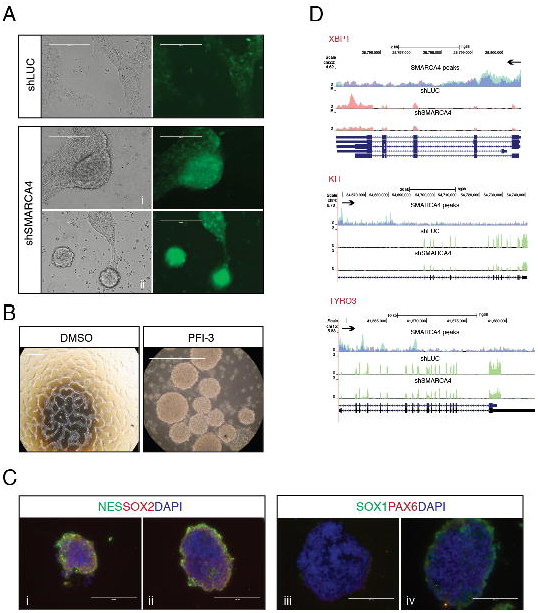
*SMARCA4* Depletion or Inhibition Causes Cell Detachment and Impedes Self-renewal. (A) Change in morphology upon shSMARCA4 treatment. The brightfield (left) and fluorescent panels (right; exhibiting the infected cells in green) are shown. Control treated cells (shLUC) grow as a monolayer (upper panel). The rounding up of the monolayer cells (i) upon SMARCA4 knockdown (shSMARCA4) followed by budding off into suspension (ii) phenotype is shown. (B) The SMARCA4 inhibitor PFI-3 phenocopies neurosphere-like formation. Images of control (DMSO) and PFI-3 treated cells at 10μM for 5 days are shown. (C) Immunofluorescence staining on cryo-sections of PFI-3 treated spheres. i-iv represent sections from different spheres. Proteins are stained by antibodies in the colors indicated. DAPI staining is shown in blue. Scale bar: 200 μm (A-B); 100 μm (C). (D) Downregulation of adhesion-related genes upon *SMARCA4* knockdown. Comparative depiction of the SMARCA4 binding (top) and the gene expression levels in the control (middle) vs knockdown (bottom) sample at the respective locus. SMARCA4 binding peaks are depicted in light blue for the eluted sample and in dark blue for the control (input) sample. Black arrows indicate the direction of transcription. SMARCA4 binding and mRNA levels are shown for *XBP1, KIT, TYRO3.* Note the binding of SMARCA4, particularly to TSS and the reduced expression of the genes upon *SMARCA4* depletion.

In order to decipher *SMARCA4’s* role in NSC self-renewal and differentiation, we opted for examining the spheres formed by PFI-3 treatment due to its general affectivity compared to transducing only a proportion of cells. Staining of spherical sections of various sizes revealed a reduction or complete loss of NSC-specific marker expression (Figure 5C); suggesting loss of stemness. Collectively, sphere formation and loss of self-renewal explain *SMARCA4* depletion in the NSC RNAi screen.

### *SMARCA4* Loss Induced Down-regulation of Adherence and Neural Suppressor Genes Can be Reversed by *SMARCA4* Overexpression

SMARCA4 has been shown to act both as a transcriptional activator as well as a repressor (Attanasio et al., 2014). Because we observed the budding off phenotype, we hypothesized that cell adherence genes might be regulated by SMARCA4. Indeed, inspection of RNA-Seq and ChIP-Seq data performed on control and *SMARCA4* knockdown NSCs revealed that expression of several adherence genes are prominently changing upon *SMARCA4* depletion. Expression of these genes, such as *XBP1, KIT*, and *TYRO3* (Figure 5D) was downregulated upon *SMARCA4* knockdown, which possibly contributes to sphere formation. This finding also supports the role of SMARCA4 in extra-cellular matrix composition, in line with two previous reports (Barutcu et al., 2016; Saladi et al., 2010).

Genome-wide ChIP-seq analysis revealed that SMARCA4 tends to bind in the vicinity of transcription start sites (TSS) (Figure 6A), and integrative analyses of the ChIP-Seq with the RNA-Seq data suggests that SMARCA4, at TSS, acts both as a transcriptional activator as well as a repressor, supporting a previous study (Attanasio et al., 2014) (Figure 6B).

**Figure 6.**
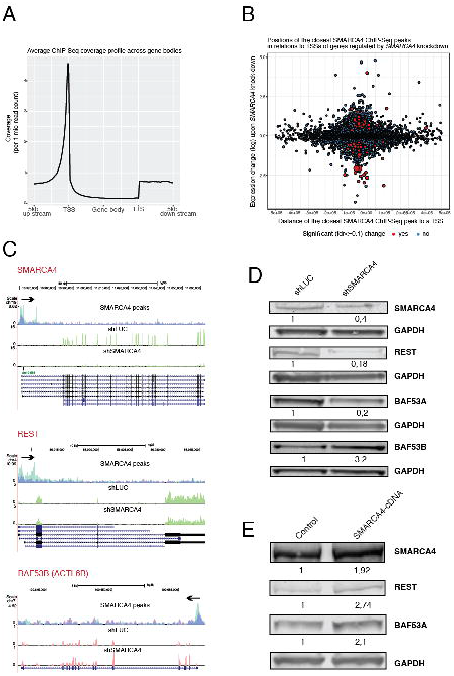
SMARCA4 de-represses neuronal genes. (A) SMARCA4 binding across genome spanning transcription start (TSS) and termination (TTS) sites, including 5kb up- and down-stream regions in NSCs. (B) Positions of the SMARCA4 binding sites (peaks) in relation to TSS of genes regulated by SMARCA4. Expression change upon knockdown (y-axis) vs distance of the closest SMARCA4 peak (x-axis) is depicted. Significant expression change is shown in red (blue: not significant). (C) Comparative depiction of the SMARCA4 binding (top) and the gene expression levels in the control (middle) vs. SMARCA4 knockdown (bottom) samples at the respective locus. SMARCA4 binding peaks are depicted in light blue for the eluted sample and in dark blue for the control (input) sample. Black arrows indicate the direction of transcription. SMARCA4 binding and mRNA levels are depicted for *SMARCA4, REST*, and *ACTL6B (BAF53B).* Note the reduced and increased expression of *REST* and *BAF53B* upon *SMARCA4* depletion, respectively. (D) Protein levels of SMARCA4, REST, BAF53A and BAF53B upon *SMARCA4* knockdown. Protein levels were normalized to GAPDH and shLUC control. Quantifications, normalized to the shLUC control are presented below the gels. (E) Protein levels of SMARCA4, REST and BAF53A upon *SMARCA4* overexpression. Protein levels were normalized to GAPDH and the control sample. Relative band intensity values are quantified below the western blots.

To investigate the depletion phenotype further, we assessed the differentiation capacity of NSC upon *SMARCA4* knockdown by looking closely at the differentially expressed genes that are regulated by SMARCA4 and are involved in NSC self-renewal or differentiation. Among the promoters SMARCA4 binds to, we identified *RE-1 silencing transcription factor* (*REST*), a repressor of neuronal differentiation, and *ACTL6B* (*BRG-1 associated factor 53, BAF53*). The BAF complex plays an important role during neural differentiation (Narayanan and Tuoc, 2014) and certain subunits are exchanged during differentiation. BAF53A is present in the neural progenitor BAF (npBAF) complex, whereas BAF53B exists in the neural BAF (nBAF) version (Lessard et al., 2007). RNA-seq data show that expression of *REST* was downregulated while *ACTL6B* was upregulated upon *SMARCA4* depletion (Figure 6C). These findings were confirmed on protein level and therefore, indicate that *SMARCA4* loss leads to a switch from BAF53A to BAF53B (Figure 6D). These results also argue that SMARCA4, together with REST, acts as a suppressor of neural differentiation, and its downregulation result in exit from the self-renewing state with skewed neural differentiation.

However, these results do not answer whether loss of self-renewal is a downstream artifact of cell-detachment followed by sphere formation or if it is directly regulated by *SMARCA4*. To this end, we aimed to investigate how expression of REST and BAF53A change upon *SMARCA4* overexpression. Interestingly, *SMARCA4* overexpression resulted in upregulation of REST by 2,7-fold and BAF53A by 2,1-fold (Figure 6E), suggesting that *SMARCA4* exerts a more direct role in stemness regulation through suppression of neuronal differentiation genes. Together, these data suggest that *SMARCA4* directly regulates NSC self-renewal by suppressing neural differentiation together with REST. Furthermore, because *SMARCA4* regulates cell adherence and NSC self-renewal independently of each other, these findings suggest that *SMARCA4* is a direct regulator of stemness in NSC.

## DISCUSSION

In humans, it has been challenging to study isogenic stem cells. The emergence of iPSC and protocols deriving other cell types from them (Zhao et al., 2013), including somatic stem cells (Sabapathy and Kumar, 2016; Sugimura et al., 2017) have opened the possibility to investigate human isogenic stem cells *ex vivo*. In this study, we exploited these advances to phenotypically compare isogenic human stem cells, and to investigate common and specific factors employed in stem cell homeostasis and differentiation in HSPCs and NSCs. Because other cell types can be derived from iPSCs by directed differentiation, we envision that this approach can be extended to other cell types.

Despite its functions are not well understood in HSPC, *SMARCA4* has been shown to be important in hematopoiesis by regulating myeloid differentiation (Holloway et al., 2003; Vradii et al., 2005), as well as in B and T cell development in the lymphoid lineage (Bossen et al., 2015; Chi et al., 2002; Choi et al., 2012). NSCs have not been studied in humans to the same extent as in model organisms: *Smarca4* loss in mouse has been shown to result in less number of NSC due to downregulation of self-renewal, and early neuronal differentiation before the onset of gliogenesis (Matsumoto et al., 2006). On the other hand, SMARCA4 is well characterized in hESC, and was shown to be a direct regulator of pluripotency (Ho et al., 2009; Zhang et al., 2014). Therefore, in our study we focused on characterizing *SMARCA4* in depth in these two adult stem cell types, HSPC and NSC.

HSPC set the starting point of our study, which we derived from easily accessible PB. For practical reasons, we used the widely applied marker CD34 to enrich for this cell population. However, sorting for CD34 is not sufficient to separate hematopoietic stem cells from progenitors. Intriguingly, we were able to distinguish these two populations during our validation experiments. After having validated the enrichment phenotype by showing the expansion of CD34^+^ population (increased cell number also in LTC-IC assay), a closer look at different hematopoietic subsets (HSC, MPP, CLP, CMP) based on immune-phenotyping revealed that HSC as well as MPP are actually depleted upon *SMARCA4* knockdown; whereas CLP and CMP populations are enriched. *In vivo* data correlated with this finding, reflected by the reduced engraftment due to impaired self-renewal of LT-HSC. These results indicate that the CD34^+^ enrichment phenotype was due to an increase in the primed-progenitor population, and not in the stem cells.

To specifically screen for genes that more directly impact HSC behavior, one could employ additional markers during the sorting process, such as CD90, CD49f, and EMCN (endomucin) (Knapp et al., 2018; Notta et al., 2011; Reckzeh et al., 2018; Wisniewski et al., 2011). However, it might be difficult to obtain enough cells to ensure full coverage of large shRNA libraries. Reducing the library size by restricting the gene count or by only employing previously validated silencing triggers might make this experiment feasible.

An interesting observation was that in the CFU assay, assessing the clonogenic potential into myeloid lineage, *SMARCA4* knockdown sample showed a 3-fold reduction in total colony numbers. Moreover, all emerging colonies were of the same type: CFU-GM. Supporting an earlier report, no erythroid colony formation was observed (Lee et al., 1999). These results suggest that *SMARCA4* knockdown traps CMP at the progenitor state and causes partial myeloid lineage specification (Holloway et al., 2003; Vradii et al., 2006). Future studies should address at which stage *SMARCA4* blocks differentiation from CMP to specified myeloid cells. Last but not least, it would be worth investigating *SMARCA4* functions in lymphoid lineage and whether a similar blockage occurs in CLP as well.

Role of SMARCA4 has been described in various mechanisms in hNSC, including cell adherence (Barutcu et al., 2016; Saladi et al., 2010). In addition, a regulatory role in transcription has also been described for SMARCA4: SMARCA4 binds and acts synergistically with REST, a zinc-finger transcription factor. REST represses its target genes by recruiting its corepressors. Its reduced activity results in expression of neural genes, implying SMARCA4 is involved in repressing neuronal-specific genes (Ooi et al., 2006). Our results reveal that the promoter of *REST* is bound by SMARCA4 and REST expression gets downregulated upon *SMARCA4* knockdown. Hence, SMARCA4 might directly regulate the expression of REST to repress neuronal differentiation. Downregulation of BAF53A and concomitant upregulation of BAF53B provide evidence for exit from self-renewal and commitment to the neural lineage. cDNA-mediated overexpression elucidates a direct link for SMARCA4’s involvement in self-renewal maintenance and indicate that neuronal differentiation upon *SMARCA4* knockdown is not an artifact of cell detachment. Last but not least, it would be interesting to investigate the interaction partners of the SMARCA4-REST complex and their target genes.

Prior studies in hESC have shown SMARCA4 to be the only catalytic subunit of a specific SWI/SNF complex, namely ESC-BAF (esBAF) (Ho et al., 2009). This complex has a unique combinatorial assembly and harbors BAF60B, which only exists within esBAF. In this study, we have not addressed the composition of the BAF complex in hiPSC. Thus, it would be interesting to investigate the BAF complex composition in iPSC, to evaluate the degree of similarity to esBAF, and its adaptation upon differentiation. Therefore, future studies aiming to gain a deeper understanding of transcriptional modifications due to combinatorial BAF subunit assembly would be of interest.

In summary, we found that *SMARCA4* not only regulates self-renewal in isogenic HSPC and NSC, but also ensures a balanced lineage specification, in a fashion unique to the cell type. SMARCA4, as a chromatin remodeler involved in numerous regulations as a transcriptional activator and repressor, poses an interesting candidate for deciphering the balance between self-renewal and differentiation at a broad range of stem cells. Finally, we show that comparative RNAi screens on isogenic human stem cells are feasible and hence, a promising approach to identify regulators of stemness and differentiation.

## EXPERIMENTAL PROCEDURES

The list of all antibodies used in this study are listed in Table S3 and primer sequences can be found in Tables S4-6.

### Cell culture

All of the primary cells and cell lines used in the study were cultivated at 37°C and 5% CO_2_ in a HERAcell 240i Incubator (Thermo Scientific).

#### HSPC

PB samples were obtained from G-CSF treated donors with their consent, and used in accordance with the guidelines approved by the Ethics Committee of the TU Dresden. Prior to CD34 enrichment, whole blood was lysed with ACK lysis buffer (Invitrogen), and enriched for CD34 by the CD34 MicroBead Kit (Miltenyi) according to the manufacturer’s instructions. For higher purity, cells were applied 2x to the magnetic column and were cultivated in StemSpan SFEM (STEMCELL) medium, supplemented with 1 uM UM729 (STEMCELL), 100 ng/ml FLT3, 100 ng/ml SCF, 50 ng/ml TPO (all R&D Systems), 100 U/ml Penicillin, 100 U/Streptomycin, and 2 mM L-glutamine (both Invitrogen). Cells were pelleted in a Centrifuge 5804R (Eppendorf) at 300 g to remove the old medium. Half medium changes were done every 2 days. 2 μg/ml puromycin was used for positive selection.

#### iPSC

Induced pluripotent stem cells (iPSCs) were cultivated in supplemented mTeSR1 (STEMCELL) with 100 U/ml Penicillin, and 100 U/Streptomycin. Fresh medium was supplied every day.

#### NSC

NSCs were cultured as previously described (Reinhardt et al., 2013). For the RNAi screen validation and further experiments involving shRNA transduction, 0.4 μg/ml puromycin was used for positive selection.

### PFI-3 treatment

The SMARCA4 inhibitor PFI-3 (Sigma-Aldrich) was added to the culture medium at a concentration of 10 μM.

### IF staining of PFI-3 treated NSC

NSCs were treated with 10 μM PFI-3 or DMSO control for 5 days. Neurospheres were pelleted for 1 min at 300 g, fixed with 4% PFA for 10 min at room temperature. Fixed samples were pelleted at 8,000 rpm for 15 sec, embedded in 300-500 μl tissue embedding medium, OCT (Slee Medical) overnight at 4°C. Cryosectioning was performed in 8 μm thick sections with an NX70 cryostat (Thermo Scientific) at −19°C. Sections were IF stained with the hNSC immunocytochemistry kit (Thermo Scientific) according to the manufacturer’s instructions. Images were acquired with an EVOS FL fluorescence microscope (Thermo Scientific) and analyzed using ImageJ image processing software.

### Lentiviral pooled shRNA screens

A custom-made DECIPHER shRNA library (Cellecta) was used for all the RNAi screens. To reach 150-fold coverage 5 million of CD34^+^ HSPCs and NSCs were transduced at an MOI of 7.5 and 0.1, respectively. To select for the shRNA carrying stem cell population, cells were sorted, at each time point, based on the shRNA reporter marker combined with CD34 (HSPC) expression (95% of NSCs were NESTIN^+^SOX2^+^, therefore, not sorted). PCR amplified gDNA carrying the shRNA barcodes was sequenced on a HiSeq2500 Illumina sequencer. Samples were 75 bp single-end sequenced in packages of 8 samples per lane, at a depth of 30 million reads. Obtained counts of reads per shRNA were converted, for each sample individually, to logarithms of the odds ratios. Subsequently, they were converted into standard scores (z-scores), where for centring and scaling, means and standard deviations of log odd ratios associated with shRNAs targeting LUC control were used. To combine z-scores associated with shRNAs targeting the same gene, a trimmed mean was calculated with 15% as a trimming factor. In case of genes targeted by 12 shRNAs, 2 most extreme results were removed from each end prior to calculating a mean z-score.

### Lentiviral transductions for validation

HSPCs were transduced as described previously (Camgoz et al., 2018), at an MOI range of 7.5-10. Except for retronectin coating, NSCs were transduced at an MOI of 0.5. Transduced cells were further cultured or analyzed by flow cytometry based on GFP expression.

### Plasmids

shRNA sequences were cloned and expressed as previously described (Camgoz et al., 2018). Constructs were confirmed by DNA sequencing.

### Transfection

NSC were transfected with the SMARCA4 expression construct pcDNA6.2/N-EmGFP-DEST (Addgene #65391) using the C-13 program with Basic Primary Neurons Nucleofector Kit (Lonza) at a Nucleofector 2b device (Lonza).

### Quantitative real time PCR (qRT-PCR)

mRNA extraction was done as described previously (Camgoz et al., 2018). Quantitative PCR was performed using the Absolute qPCR 2x SYBR Green Kit (Thermo Scientific) following the manufacturer’s protocol on a CFX96 Real-Time System (Bio-Rad). Using the 2-ΔΔCt method, mRNA levels were normalized to the LUC control against GAPDH as an internal control.

### Western blot

Samples were processed, loaded on SDS gels and transferred as previously described (Camgoz et al., 2018). Typically, 1 million cells were lysed in 40 μl 1 x cell lysis buffer (Cell Signaling) supplemented with 1x RIPA lysis buffer (Santa Cruz). Bands were visualized on a LI-COR Odyssey imaging system using the intensity 3.5 for 680LT dye and intensity 7.0 for 800CW dye, and were analyzed using Image Studio Lite (Version4, LI-COR).

### Long-term culture-initiating cell assay (LTC-IC)

For the LTC-IC assay CD34^+^ HSPCs were transduced with *SMARCA4* and control shRNAs, and were subjected to 2 μg/ml puromycin selection between 2-4 dpt. 1000 cells/well were seeded on irradiated M2-10B4 murine bone marrow stromal cells in a 96-well plate and cultured in Myelocult H5100 (STEMCELL) according to the manufacturer’s instructions (n=5). Between week 2-5, cobblestones, defined as an area of at least 5 tightly adjacent hematopoietic cells with a rectangular shape, were manually counted using a Celigo cytometer (Brooks).

### Colony forming unit assay (CFU)

CD34^+^ HSPC were transduced with *SMARCA4* and control shRNAs, and subjected to 2 μg/ml puromycin selection for 2 days. 1000 cells /dish were seeded in 3 replicates in 35-mm dishes with methylcellulose-containing medium supplemented with rhSCF, rhGM-CSF, rhG-CSF, rhIL-3, rhIL-6, and rhEPO (MethoCult H4435 Enriched; STEMCELL). After 14 days colony types and numbers were analyzed using a STEMvision™ instrument (STEMCELL).

### Mice

NOD.Cg-*Prkdc^scid^ Il2rg^tm1Wjl^*/SzJ *Kit^W41/W41^* mice (NSGW41) were generated by backcrossing the *Kit^W41^* allele (C57BL/6-Kit < W-41 >) to NSG mice for 16 generations. The resulting heterozygous NSG *Kit^W41/+^* mice were intercrossed to receive the homozygous NSGW41 mice used for experiments (Cosgun et al., 2014). All mice were bred and maintained in individually ventilated cages under specific pathogen-free conditions at the experimental center of the TU Dresden. The Landesdirektion Dresden, as responsible authority, approved all animal experiments.

### Transplantation

For humanization 5×10^5^ CD34-enriched cells were injected intravenously in 150ul PBS/ 5 % FCS into 7- to 11-week-old unconditioned NSGW41 mice. After transplantation mice were given neomycin-containing drinking water for 3 weeks.

### Flow cytometry analysis and cell sorting

BM and blood samples were collected and prepared as described before (Cosgun et al., 2014). Samples were acquired on a LSRII cytometer (BD Biosciences).

All the other antibody stainings were performed according to the manufacturer’s instructions. Samples were acquired in PBS-EB (0.5M EDTA ^+^ 0.4% [m/v] BSA in PBS) on a MACS Quant Analyzer (Miltenyi), FACS Aria III sorter (BD Biosciences) or FACS Canto II SORP (BD Biosciences). Data were analyzed using FlowJo software (TreeStar).

### Chromatin immunoprecipitation sequencing (ChIP-seq)

Chromatin immunoprecipitation was done as described previously (Ding et al., 2015) using 20 μl of ChIP-grade SMARCA4. DNA was purified and eluted using a PCR purification kit (Qiagen) in 30 μl HPLC-grade water. Samples were submitted to NGS on an Illumina HiSeq 2000. Gene expression levels were estimated with kallisto software (ver. 0.43.0), using cDNA sequences from Ensembl database (release 79) as a reference, and further differential expression analysis was performed using sleuth algorithm (ver. 0.28.1).

### RNA-seq

RNA samples were harvested in parallel to gDNA isolation at each time point during the RNAi screens. In addition, untreated iPSC and NSC RNA samples were sequenced as part of their characterization. RNA samples were sequenced with poly-dT enrichment using single-end sequencing. Each sample was sequenced at a depth of yielding at least 20 million reads.

### Data and statistical analysis

Data were analyzed using GraphPad Prism version 6 (GraphPad Software). Results were presented as standard error of the mean (SEM, presented as error bars). Comparisons between experimental groups were made using unpaired Student’s two-tailed *t* test. p < 0.05 was considered to be statistically significant.

## ACCESSION NUMBERS

Deep sequencing datasets of RNA-seq and ChIP-seq experiments have been submitted to GEO and the accession numbers will soon be provided upon approval.

## AUTHOR CONTRIBUTION

C.G. planned the research and executed the experiments; S.R. and C.W. performed the transplantation and bone marrow analysis; C.G. and M.P.-R. analyzed the data; S.K. performed the iPSC characterization experiments; M.W. and M.B. contributed materials; A.D. performed the deep-sequencing experiments; C.G. and F.B. wrote the manuscript with input from all the authors.

## ACKNOWLEDGEMENTS

We would like to thank Katja Bernhardt and Anne Gompf from the Core Flow Cytometry for their help with the FACS analysis. The excellent help of Srimasorn Sumitra during NSC derivation is greatly acknowledged. We also thank all Buchholz laboratory members for their valuable feedback and discussions. The project was supported by grants from Deutsche Krebshilfe (SyTASC / 70111969) and the DFG SFB655 (B5). C.W. was supported by the German Research Foundation (DFG) through FOR2033-A03, TRR127-A5, WA2837/6-1 and WA2837/7-1.

## Tables

Table S2. shRNA read counts and z-scores from the HSC- and NSC-RNAi screens (excel file)

