## Supplemental Information for "Comparative RNAi Screens in Isogenic Human Stem Cells Reveal *SMARCA4* as a Differential Regulator"

**Figure S1.** Related to Figure 2.

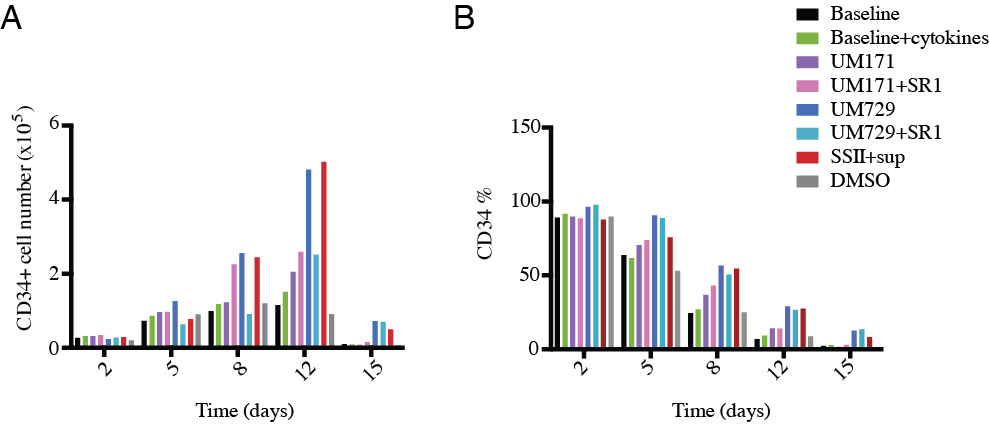

**Figure S1. Optimization of the CD34 HSPC culture conditions** Comparison of various suspension culture conditions to evaluate **(A)** CD34^+^ cell numbers as well as **(B)** percentage, indicating the proliferation vs differentiation, respectively. SR1: StemRegenin1; SSII: StemSpan SFEM II; sup: CD34^+^ Expansion Supplement.

**Figure S2.** Related to Figure 2.

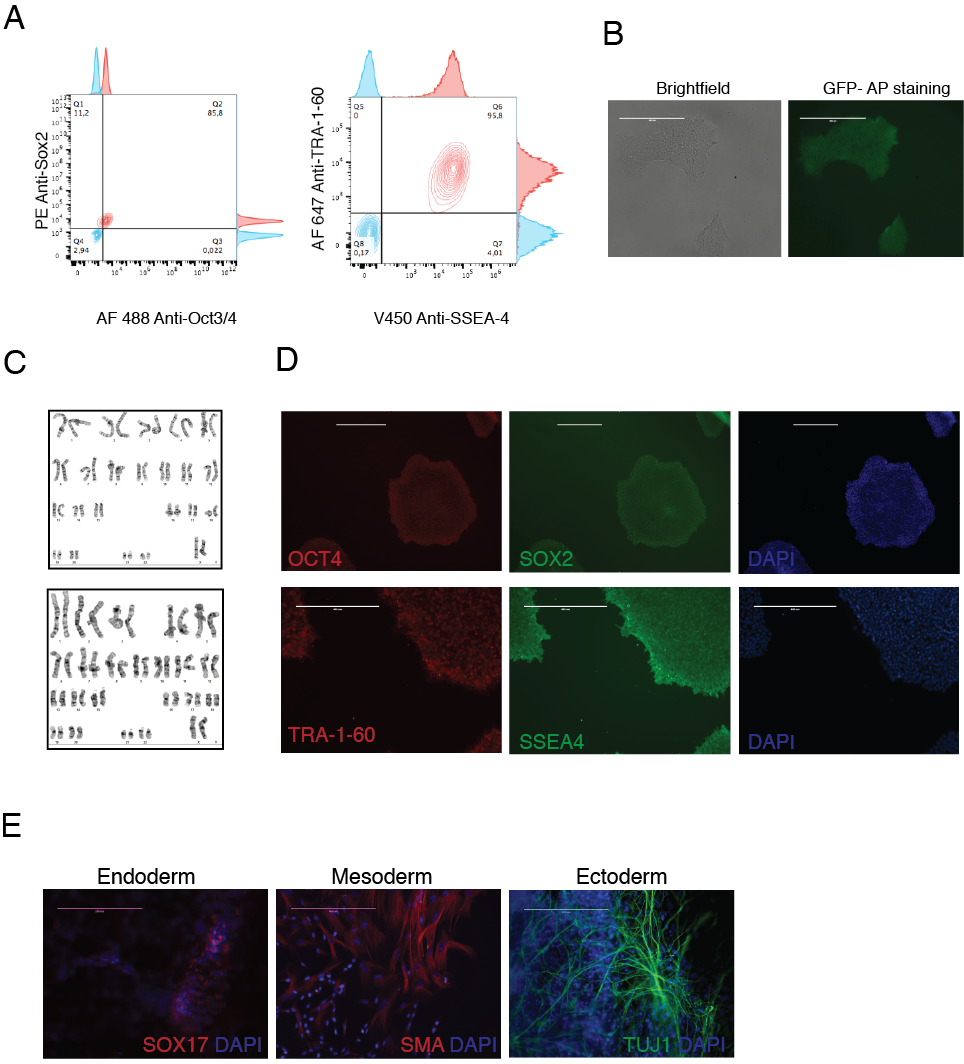

**Figure S2. Characterization of the HSC-derived iPSC lines** **(A)** Flow cytometry analysis pluripotency markers. **(B)** GFP-coupled alkaline phosphatase (AP) staining. **(C)** G-band karyotyping of the HSC-derived two iPSC lines. **(D)** Immunofluorescence staining of nuclear (OCT4 and SOX2) and surface (TRA-1-60 and SSEA-4) iPSC markers. **(E)** Immunofluorescence staining upon differentiation into the 3 germ-layers. Scale bar: 400 μm (Endoderm: 200 μm).

**Figure S3.** Related to Figure 2.

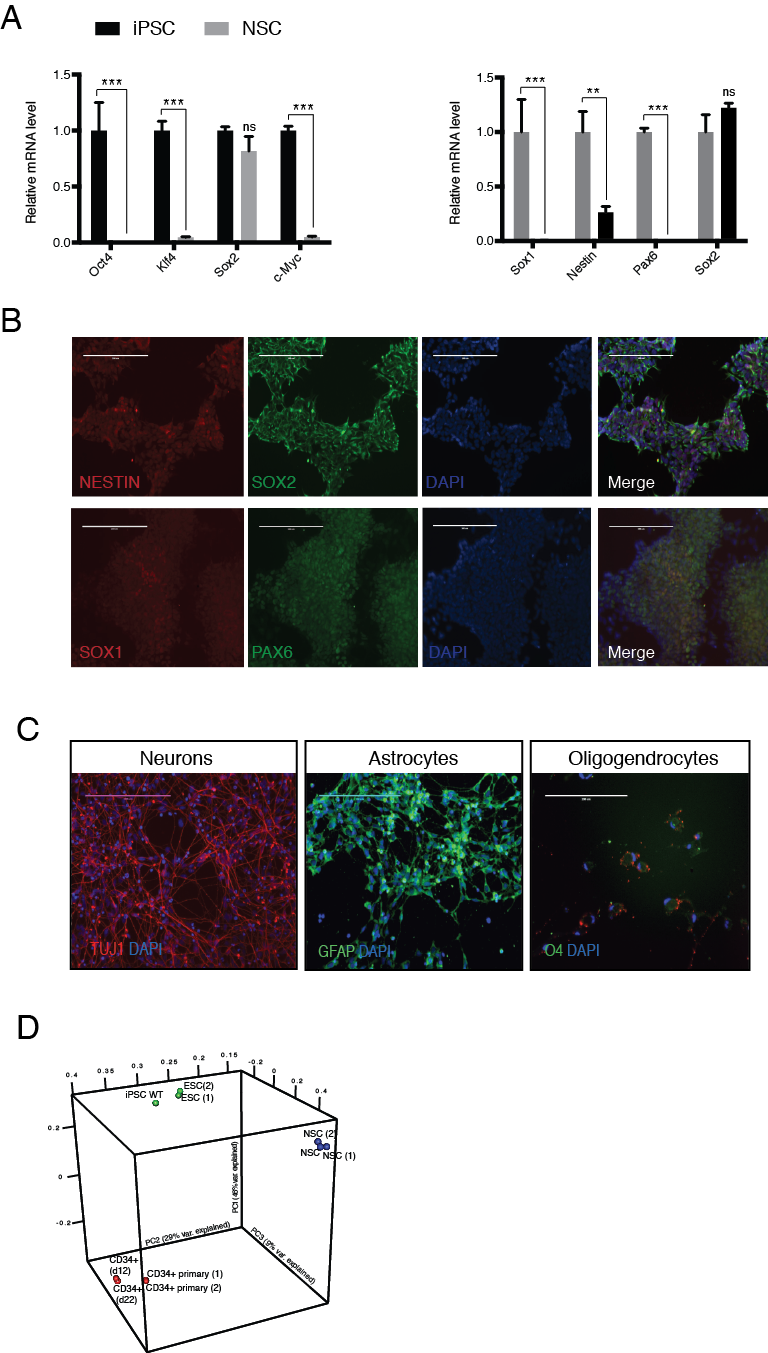

**Figure S3. Characterization of the iPSC-derived NSC lines** **(A)** Change in gene expression upon neural induction of iPSC measured by qRT-PCR. Data are represented as mean ± SEM. All expression changes, except for *SOX2,* are statistically significant. **(B)** Immunofluorescence staining of nuclear (SOX1, SOX2, PAX6) and cytoplasmic (NESTIN) NSC markers. **(C)** Immunofluorescence staining of differentiated NSC into neurons (TUJ1), astrocytes (GFAP), and oligodendrocytes (O4). **(D)** Principal Component Analysis (PCA) cluster each isogenic cell types to its primary and cell line counterpart from literature. Scale bar: 200 μm. **p < 0.01, ***p < 0.001; ns, not significant.

**Table S1. STR analysis of the isogenic HSC, iPSC, and NSC.** Related to Figure 1. Values displayed for each cell type shows the number of repeats of a DNA fragment at a length of 2-13 bp, to which probes were attached and PCR amplified.

| **Fragment**  **Cell type** | **HSC** | **iPSC #1** | **iPSC #2** | **NSC #1** | **NSC #2** |
| --- | --- | --- | --- | --- | --- |
| Amelogenin | X | X | X | X | X |
| THO1 | 6, 9.3 | 6, 9.3 | 6, 9.3 | 6, 9.3 | 6, 9.3 |
| D3S1358 | 16, 17 | 16, 17 | 16, 17 | 16, 17 | 16, 17 |
| VWA | 16, 19 | 16, 19 | 16, 19 | 16, 19 | 16, 19 |
| D21S11 | 30, 31.2 | 30, 31.2 | 30, 31.2 | 30, 31.2 | 30, 31.2 |
| TPOX | 8, 11 | 8, 11 | 8, 11 | 8, 11 | 8, 11 |
| D7S820 | 8, 11 | 8, 11 | 8, 11 | 8, 11 | 8, 11 |
| D19S433 | 14 | 14 | 14 | 14 | 14 |
| D5S818 | 8, 12 | 8, 12 | 8, 12 | 8, 12 | 8, 12 |
| D2S1338 | 23, 25 | 23, 25 | 23, 25 | 23, 25 | 23, 25 |
| D16S539 | 9, 13 | 9, 13 | 9, 13 | 9, 13 | 9, 13 |
| CSF1PO | 10, 11 | 10, 11 | 10, 11 | 10, 11 | 10, 11 |
| D13S317 | 12, 13 | 12, 13 | 12, 13 | 12, 13 | 12, 13 |
| FGA | 20, 26 | 20, 26 | 20, 26 | 20, 26 | 20, 26 |
| D18S51 | 16, 21 | 16, 21 | 16, 21 | 16, 21 | 16, 21 |
| D8S1179 | 12, 15 | 12, 15 | 12, 15 | 12, 15 | 12, 15 |

Table S3. Summary of antibodies used in this study. Related to Figures 2-6.

| **Target** | **Clone ID** | **Provider** |
| --- | --- | --- |
| CD34 | 8G12 | BD Biosciences |
| CD38 | 60014AZ | Stemcell Technologies |
| CD45RA | 560673 | BD Biosciences |
| hCD45 | HI30-PB | Biolegend |
| mCD45 | 30F11-A780 | eBioscience |
| CD49f | 60037PB | Stemcell Technologies |
| CD90 | 5E10 | Stemcell Technologies |
| TRA-1-60 | 560850 | BD Biosciences |
| SSEA4 | 561156 | BD Biosciences |
| OCT3/4 | 560253 | BD Biosciences |
| SOX2 | 560291 and 245610 | BD Biosciences |
| NESTIN | Clone 25/NESTIN | BD Biosciences |
| SOX17 | ab84990 | Abcam |
| SMA | 180106 | Life Technologies |
| TUJ1 | A25532 | Life Technologies |
| GFAP | GA5 | Merck |
| O4 | 81 | Merck |
| SMARCA4 (ChIP grade) | EPNCIR111A | Abcam |
| REST | ab21635 | Abcam |
| BAF53A | ab174941 | Abcam |
| BAF53B | PA5-25247 | Thermo Scientific |
| GAPDH | AP16240PU-N | Acris |
| IRDye 680LT | 925-68023 | LI-COR |
| IRDye 800 CW | 925-32210 | LI-COR |
| Pluripotent Stem Cell 4-Marker Immunocytochemistry kit | A24881 | Thermo Scientific |
| Human Neural Stem Cell Immunocytochemistry kit | A24354 | Thermo Scientific |
| 3-Germ Layer Immunocytochemistry kit | A25538 | Thermo Scientific |
| FOXA2 | MAB2400-100 | R&D Systems |
| NCAM | ab75813 | Abcam |
| T-Brachyury | ab140661 | Abcam |
| PAX6 | ab5790 | Abcam |
| NESTIN | ab6320 | Abcam |

**Table S4. Primers sequences used in nested PCR for gDNA amplification.** Related to Figure 2. NF2 sequence in Gex1-NF2 primers is shown in red. Barcode sequences special for each MLPX primer are indicated in bold.

| **Name of the primer** | **Forward primer (5^’^-3^’^)** |
| --- | --- |
| F2 | TCGGATTCGCACCAGCACGCTA |
| GexSeqS | AGAGGTTCAGAGTTCTACAGTCCGAA |
| Gex1-NF2 | TCAAGCAGAAGACGGCATACGATCGCACCAGCACGCTACGCA |
| MLPX 45 | **ACATCG**AGAGGTTCAGAGTTC**TACAGTCCGAA** |
| MLPX 47 | **TGGTCA**AGAGGTTCAGAGTTC**TACAGTCCGAA** |
| MLPX 49 | **ATTGGC**AGAGGTTCAGAGTTC |
| MLPX 50 | **GATCTG**AGAGGTTCAGAGTTC |
| MLPX 51 | **TCAAGT**AGAGGTTCAGAGTTC |
| MLPX 53 | **AAGCTA**AGAGGTTCAGAGTTC |
| MLPX 54 | **GTAGCC**AGAGGTTCAGAGTTC |
| MLPX 55 | **TACAAG**AGAGGTTCAGAGTTC |

**Table S5. Primer sequences used for cloning the RNAi hits for further validation.** Related to Figure 2.

| **Gene** | **Forward primer (5’-3’)** | **Reverse primer (5’-3’)** |
| --- | --- | --- |
| *SMARCA4* | ACCGGGCCAAGTAAGATGTTGATGATGTTAATATTCATAGCATCATCGACATCTTGCTTGGCTTTT | CGAAAAAAGCCAAGCAAGATGTCGATGATGCTATGAATATTAACATCATCAACATCTTACTTGGCC |
| *LUC* | ACCGGCTTCGAAATGTTTGTTTGGTTGTTAATATTCATAGCAACCAAACGAACATTTCGAAGTTTT | CGAAAAAACTTCGAAATGTTCGTTTGGTTGCTATGAATATTAACAACCAAACAAACATTTCGAAGC |

**Table S6. Primer sequences used for qRT-PCR.** Related to Figure 2.

| **Gene name** | **Forward primer (5’-3’)** | **Reverse primer (5’-3’)** |
| --- | --- | --- |
| *SMARCA4* | CTCGGTCCGTCAAAGTGAAGA | AGCGGTCCTCCTCTTGTTCC |
| *KLF4* | ACCCACACAGGTGAGAAACC | ATGCTCGGTCGCATTTTTGG |
| *OCT4* | GGAAGGAATTGGGAACACAAAGG | AACTTCACCTTCCCTCCAACCA |
| *SOX2* | TGGCGAACCATCTCTGTGGT | CCAACGGTGTCAACCTGCAT |
| *c-MYC* | CCAGCAGCGACTCTGAGGA | GAGCCTGCCTCTTTTCCACAG |
| *SOX1* | ATACTGGAGACGAACGCCG | AACCCAAGTCTGGTGTCAGC |
| *PAX6* | CCAGCCAGACCTCCTCATAC | TGGCTGACTGTTCATGTGTGTC |
| *NESTIN* | CTCAGCTTTCAGGACCCCAA | GTCTCAAGGGTAGCAGGCAA |
| *GAPDH* | GCACCGTCAAGGCTGAGAAC | AGGGATCTCGCTCCTGGAA |

**Supplemental Experimental Procedures**

**HSC suspension culture condition optimization**

For optimizing HSC suspension culture conditions, several medium compositions were tested. The following medium recipes are given with the cytokine mixture of SCF, FLT-3, and TPO (all R&D Systems) in StemSpanSFEM (STEMCELL) unless otherwise stated, 100 U/ml Penicillin/Streptomycin, 2 mM L-glutamine, 100 ul heat-inactivated FCS (all Invitrogen). Baseline: 10 ng/ml of SCF, FLT-3, TPO. Baseline+cytokines: 100 ng/ml SCF and FLT-3, 50 ng/ml TPO. UM171: 0,35 µM UM171 (STEMCELL), 100 ng/ml SCF and FLT-3, 50 ng/ml TPO. UM171+SR1: 0,35 µM UM171, 0,1 mM SR1 (ChemieTek), 100 ng/ml SCF and FLT-3, 50 ng/ml TPO. UM729: 1 µM UM729 (STEMCELL), 100 ng/ml SCF and FLT-3, 50 ng/ml TPO. UM729+SR1: 1 µM UM729, 0,1 mM SR1, 100 ng/ml SCF and FLT-3, 50 ng/ml TPO. SSII+sup: StemSpanSFEM II and 10x CD34^+^ expansion supplement (both STEMCELL). DMSO: 1:5000 DMSO.

**Reprogramming**

2x10^5^ CD34 enriched HSPCs were cultivated for 3 days in StemSpan SFEM medium with 100 ng/ml SCF, 50 ng/ml IL-3, 25 ng/ml GM-SCF, 100 U/ml Penicillin, 100 U/Streptomycin, and 2 mM L-glutamine. On the day of transduction (day 0), cells were plated on matrigel-coated plates (Corning). Reprogramming factors were delivered by CytoTune 2.0 Sendai virus (Invitrogen) according to the manufacturer’s instructions with 4 µg/ml polybrene to improve transduction efficiency. Until day 8, Sendai virus’ manufacturer’s instructions were applied using ReproTeSR (STEMCELL). As of 8 dpt daily medium changes were carried out with 2 ml fresh ReproTeSR, as instructed by the manufacturer. Colonies were picked between 21-28 dpt, transferred to matrigel-coated plates and cultivated in mTeSR1.

**NSC derivation**

iPSC was differentiated to NSC as previously described (Reinhardt et al., 2013). 2 NSC lines were derived and characterized with IF staining and uninduced differentiation.

**IF staining for iPSC and NSC characterization**

As part of the characterization assays, iPSC and NSC were stained with their respective stem cell specific using the pluripotent stem cell 4-marker immunocytochemistry kit and human neural stem cell immunocytochemistry kit, respectively, according to the manufacturer’s instructions (Thermo Scientific). Additionally, iPSC was stained with alkaline phosphatase (AP) live stain according to the manufacturer’s instructions (Thermo Scientific).

Embryoid bodies (EBs) were derived from iPSC and differentiated spontaneously over 14 days. Differentiated EBs were stained with 3-Germ Layer immunocytochemistry kit according to the manufacturer’s instructions, for mesoderm and ectoderm markers. For endoderm staining, SOX17 antibody (abcam) was used. Cells were washed twice with 1x PBS, and fixed with 4% paraformaldehyde (PFA) for 10 min at room temperature. Cells were washed with wash buffer for 5-10 min. Afterwards, cells were blocked for 1 h with blocking solution (0.5% BSA in 0.3 Triton X-100 in PBS), and primary antibody recognizing SOX17 was diluted in blocking solution and incubated overnight at 4°C. Cells were washed 3 times with 1x PBS for 5 min each, followed by 1 h incubation with the secondary antibody (1:250 diluted in blocking solution) at room temperature. As secondary antibody AF488 goat anti-mouse antibody (1:250, Thermo Scientific) was used. After 3 final washed with 1x PBS, cells in 1x PBS were stained in nucleus with DAPI containing NucBlue solution from the 3-Germ Layer immunocytochemistry kit.

NSCs, which were spontaneously differentiated over 3 weeks, were washed once with 1x PBS, and fixed with 4% PFA for 20 min at room temperature. Afterwards, cells were permeabilized and blocked for 45 min with 10% FCS, and 1% BSA in PBS, and 0.1 Triton X-100 (PBS-T). Cells were washed once with washing buffer (0.1% BSA in PBS) and incubated overnight at 4°C with primary antibodies recognizing TUJ1, O4, and GFAP. The next day cells were washed 3 times with washing buffer, followed by secondary antibody (diluted 1:250 in 0.1% BSA in PBS-T) incubation for 1 h at room temperature. As secondary antibodies AF647 donkey anti-rabbit against for TUJ1, AF88 goat anti-mouse against O4, and AF488 donkey anti-mouse against GFAP were used. Cells were washed 3 times with washing buffer, and covered with 1x PBS. Finally, DAPI staining was done with 1 drop of NucBlue per well and cells were imaged. Images were acquired with an EVOS FL fluorescence microscope (Thermo Scientific) using a 4x, 10x, and 20x objectives. Images were analyzed using ImageJ image processing software.
