## Supplementary figures and images for "Comparative RNAi Screens in Isogenic Human Stem Cells Reveal *SMARCA4* as a Differential Regulator"

### Supplemental Figure 1

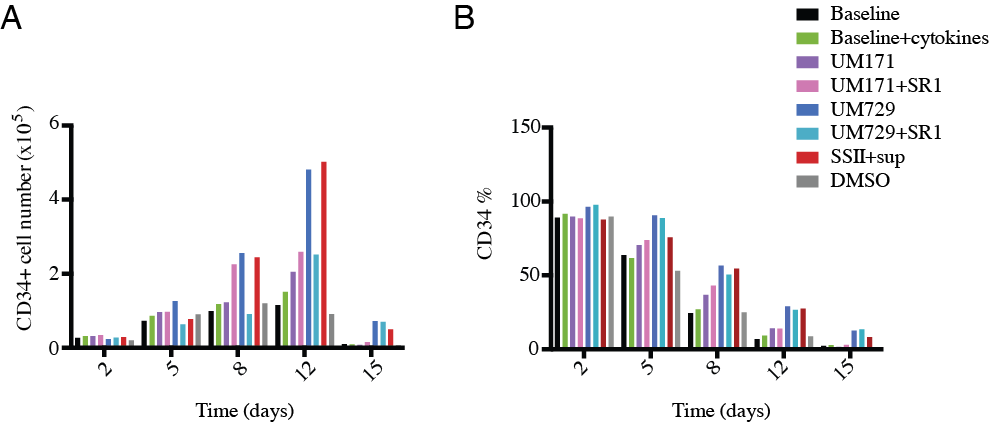

### Supplemental Figure 2

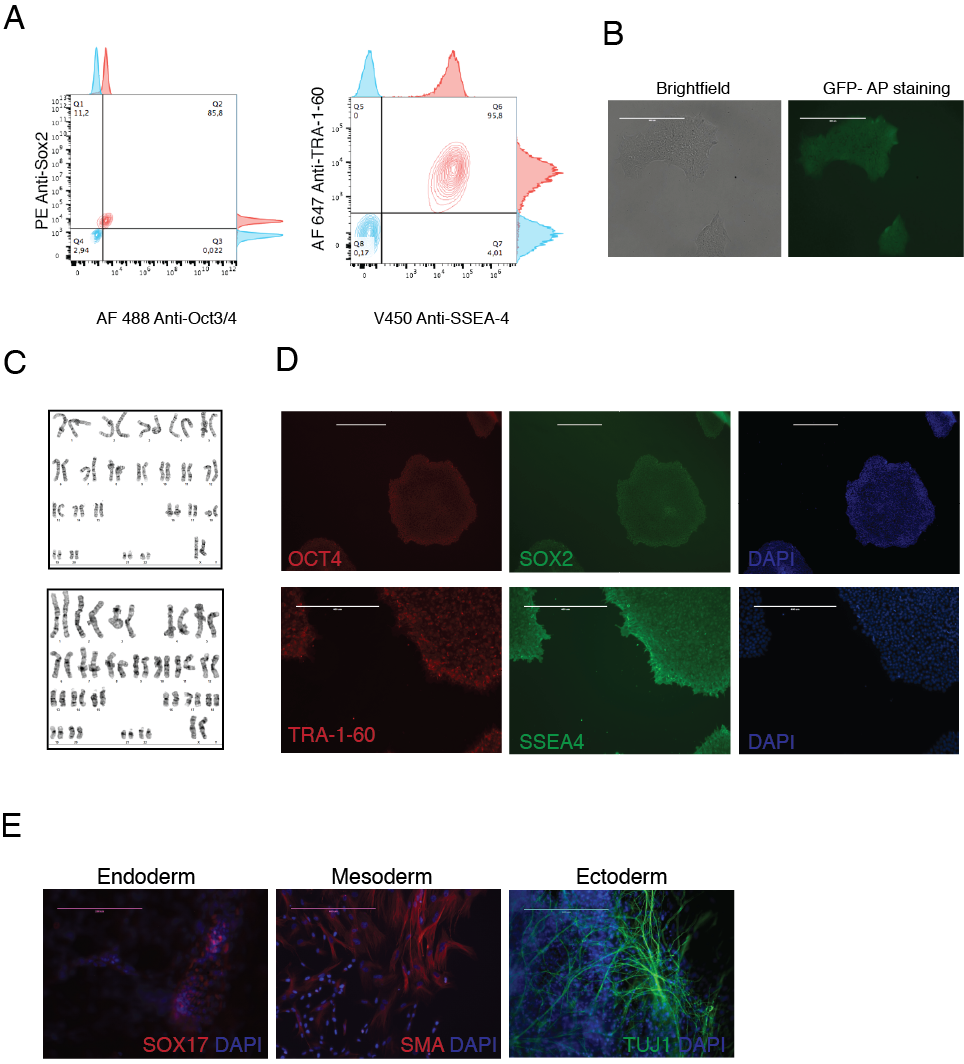

### Supplemental Figure 3

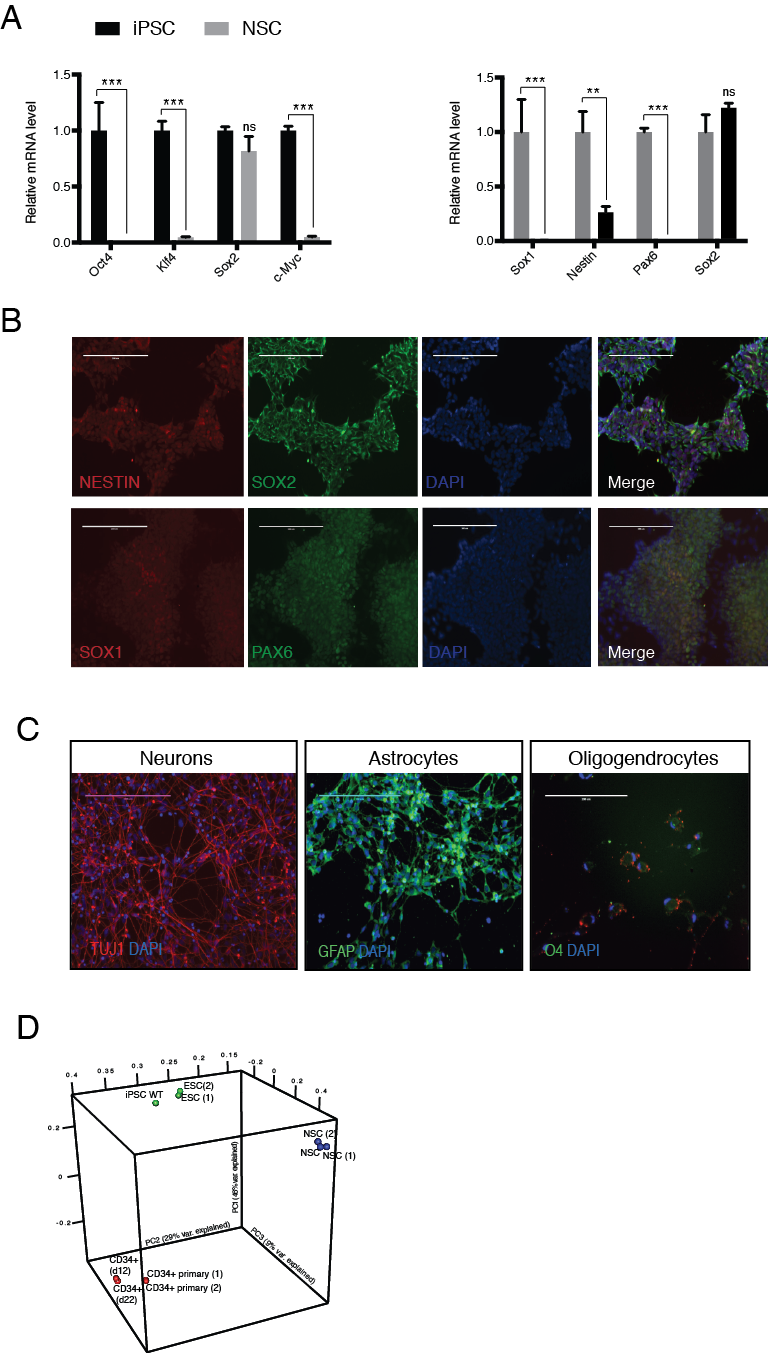
